# Cytometric Analysis of Diverse Glaucophyte Species Reveals Distinctive Signals Useful for Fluorescence-Based Detection and Sorting

**DOI:** 10.1101/2021.10.21.465165

**Authors:** Rossella Calvaruso, Janice Lawrence, Adrian Reyes-Prieto

## Abstract

Glaucophytes, red algae and viridiplants (green algae and land plants) are formally united in the supergroup Archaeplastida. Although diverse molecular and genomic evidence suggest the common origin of the three Archaeplastida lineages, the lack of a robust glaucophyte knowledgebase has limited comprehensive evaluations of competing hypotheses. Glaucophytes are rare and apparently confined to freshwater habitats. However, the distribution and diversity of these algae have not been thoroughly explored owing to challenges with detecting and isolating novel specimens. Here we examined the cytometric signatures of representative species of the genera *Cyanophora*, *Cyanoptyche*, *Glaucocystis* and *Gloeochaete* for a distinctive signal that would aid identification. Most glaucophytes analyzed presented a relatively high red fluorescence signal due to the presence of the blue phycobiliproteins C-phycocyanin and allophycocyanin. Cell-size differences and the concurrent presence of the red phycobiliprotein phycoerythrin in other algal lineages, such as red algae and cryptophytes, allowed us to distinguish glaucophytes from other photosynthetic cells containing blue phycobiliproteins. Our results indicate that the peculiar autofluorescence signal of glaucophytes will facilitate further identification and isolation on novel specimens of this scarce but important algal group.

## Introduction

Glaucophyta (Skuja 1948), Rhodophyceae (red algae) and Chloroplastida (green algae and land plants; also called Viridiplantae) are the three lineages harboring primary plastids united in the Archaeplastida (Adl et al. 2005). Early phylogenomic studies using nuclear encoded proteins resolved the Archaeplastida (Plantae) as a monophyletic group (Rodríguez-Ezpeleta et al. 2005; Hackett et al. 2007). However, posterior analyses comprising expanded sequence data sets, broader taxonomic samples and novel taxa have not recovered the Archaeplastida lineages in a single clade (Burki et al. 2012; Brown et al. 2013; Yabuki et al. 2014; Baurain et al. 2010; Burki et al. 2016; Janouškovec et al. 2017; Brown et al. 2017; Heiss et al. 2018; Gawryluk et al., 2019; Schön et al, 2021). In addition to the intrinsic limitations of the methods used for phylogenetic estimation (Burki et al. 2016; Mackiewicz and Gagat; 2014), the scarce amount of glaucophyte data, restricted in most studies to *Cyanophora paradoxa* and *Glaucocystis* spp., has been a drawback when exploring the evolutionary history of the Archaeplastida. Analyses of mitochondrial genomic data recovered the Archaeplastida clade, with weak to moderate node support, only when the glaucophyte sample was expanded to include all genera available in culture collections (Jackson and Reyes-Prieto 2014). This result demonstrates that a broader glaucophyte database is critical for robust investigations of Archaeplastida evolution (Jackson and Reyes-Prieto 2014; Mackiewicz and Gagat 2014).

Sampling evidence has suggested that glaucophytes are restricted to freshwater environments with specimens collected from the water column or benthos of rivers, lakes, fishponds and even soils. However, a more recent record from the Tara Ocean project revealed glaucophyte metabarcode sequences from oceanic samples (de Vargas et al. 2015). Benthic species are frequently attached to submerged plant leaves, detritus, or filamentous algae. Even though glaucophytes are distributed in a variety of freshwater habitats, new specimens are rarely isolated, and reports are mainly restricted to the relatively well-known genera *Cyanophora* and *Glaucocystis*. In the case of *Cyanophora*, a planktonic species, isolates have been reported from ephemeral ponds, lakes and rivers (Barone et al. 2006; Korshikov 1941; Kugrens et al. 1999; Skuja 1956). Specimens of the benthic genus *Glaucocystis* are relatively more abundant (Whitton 2002; Scholz and Liebezeit 2012; Sheath and Steinman 1982) and have been reported as epiphytes on filamentous algae, such as *Spirogyra* and *Oedogodium* (Hoffmann and Kostikov 2015). Other glaucophytes, such as *Cyanoptyche gloeocystis*, have been referred as “quite common” (Sheath and Steinman 1982; Fenwick 1966), but few records are actually available. A relatively recent report of *Chalarodora azurea* in Northern Slovakia described the cells as attached to filaments of the cyanobacterium *Hapalosiphon fontinalis* (Hindak and Hindakova 2012). Overall, studies of glaucophyte distribution and species diversity are limited, and in most cases only provide vague identifications and minimal descriptions of the habitat. As a consequence, robust examinations of the relationships among the few known genera are also rare. However, some investigations have provided novel insights into the evolution of the group (Smith et al. 2014; Jackson and Reyes-Prieto 2014; Jackson et al. 2014), revealed cryptic species diversity (Chong et al. 2014) and motivated revisions of taxonomic delimitations (Takahashi et al. 2014; Takahashi et al. 2016).

Our aim was to establish simple detection methods that increase chances of further identification and isolation of glaucophyte cells from the environment using the peculiar glaucophyte collection of photosynthetic pigments as reference to detect these algal cells in high-throughput surveys. The plastids of glaucophytes are unique among the photosynthetic organelles from other algal groups due to the conspicuous presence of the blue phycobiliproteins (antenna photopigments) C-phycocyanin (CPC) and allophycocyanin (APC) (Chapman 1966; Schmidt et al. 1979; Watanabe et al. 2012). Blue phycobiliproteins are also found in cyanobacteria and plastids of red algae and cryptophytes (Watanabe et al. 2012; MacColl 1998; Gantt and Lipschultz 1974; Beutler et al. 2004; MacColl and Berns 1978; Lichtlé et al. 1992; McKay et al. 1992), but the latter two groups usually harbor phycoerythrin (PE; a red phycobiliprotein) as the main antenna photopigment (Beutler et al. 2004; Gantt and Lipschultz 1974; Hoef-Emden 2008; Bryant 1982). In contrast, previous studies have not detected PE as part of the *Cyanophora* and *Glaucocystis* light harvesting machinery (Chapman 1966; Misumi & Sonoike 2017). We suggest that the distinctive collection of phycobiliproteins in the glaucophyte plastids is a useful trait to identify and isolate cells of this lineage using fluorescence-based detection (see Table 1).

**Table 1.**
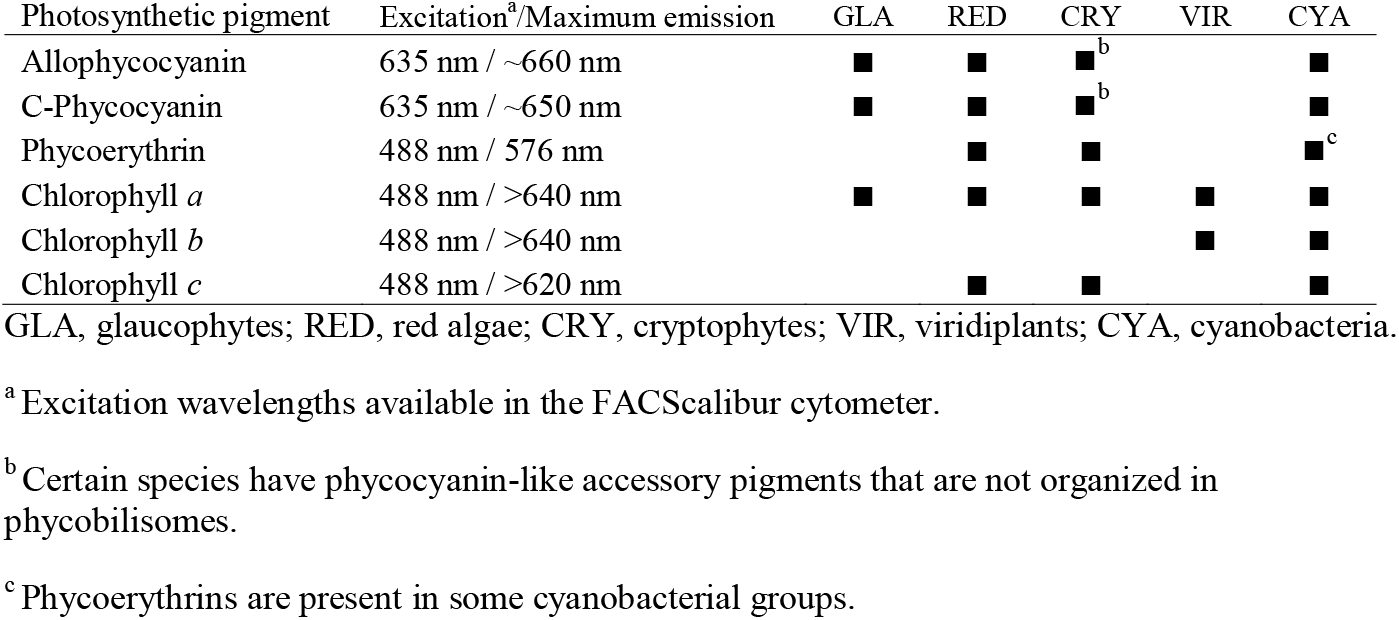
Principal photosynthetic pigments present in diverse groups of photosynthetic lineages.

| Photosynthetic pigment | Excitation<sup>a</sup>/Maximum emission | GLA | RED | CRY | VIR | CYA |
| --- | --- | --- | --- | --- | --- | --- |
| Allophycocyanin | 635 nm / ~660 nm | ■ | ■ | ■<sup>b</sup> |  | ■ |
| C-Phycocyanin | 635 nm / ~650 nm | ■ | ■ | ■<sup>b</sup> |  | ■ |
| Phycoerythrin | 488 nm / 576 nm |  | ■ | ■ |  | ■<sup>c</sup> |
| Chlorophyll *a* | 488 nm / >640 nm | ■ | ■ | ■ | ■ | ■ |
| Chlorophyll *b* | 488 nm / >640 nm |  |  |  | ■ | ■ |
| Chlorophyll *c* | 488 nm / >620 nm |  | ■ | ■ |  | ■ |

GLA, glaucophytes; RED, red algae; CRY, cryptophytes; VIR, viridiplants; CYA, cyanobacteria.
<sup>a</sup> Excitation wavelengths available in the FACScalibur cytometer.
<sup>b</sup> Certain species have phycocyanin-like accessory pigments that are not organized in phycobilisomes.
<sup>c</sup> Phycoerythrins are present in some cyanobacterial groups.

Multi-parameter flow cytometry coupled to fluorescence-activated cell sorting (FACS) has been used extensively to identify and recover particular populations of photosynthetic cells (Marie et al. 2005; Marie et al. 2010; Campbell 2001; Davey and Kell 1996; Becker et al. 2002; Zhou et al. 2012, Dunker and Wilhelm 2018; Xie et al. 2018). These methods have also been applied to ecological investigations (Cellamare et al. 2010; Sinigalliano et al. 2010, Liu et al 2018), the selection of populations with commercial relevance (e.g., algae with high lipid content; Manandhar-Shrestha and Hildebrand 2013), microbial diversity surveys (Davey and Kell 1996;

Zhou et al. 2012, Props et al. 2018) and for the establishment of axenic cultures (Sensen et al. 1993). Here we present cytometric analyses of diverse glaucophyte species, propose the use of a distinctive fluorescence signature for cell sorting, and describe a simple FACS-based method to establish cell cultures.

## Materials and methods

### Autofluorescence analysis

Representative species of the genera *Cyanophora, Cyanoptyche, Glaucocystis* and *Gloeochaete* were analyzed to produce a baseline catalogue of the glaucophyte autofluorescence signature (see Table 2). Cell cultures were maintained in liquid DY-V media (Andersen et al. 2005) supplemented with 0.5 mM urea at 18 °C on a 14/10 h light/dark cycle (20 μmol photons m^−2^ s^−1^). Aliquots (2.5 ml) from cultures were filtered using 100-μm cell strainers and then analyzed using a benchtop FACSCalibur flow cytometer (Becton Dickinson) equipped with both a blue argon-ion laser (488 nm) and a red-diode laser (635 nm). Emitted signals were collected using the forward scatter detector (FSC) along with fluorescence detectors equipped with bandpass filters of 530/30 nm (green fluorescence; FL1 detector), 580/42 nm (orange fluorescence; FL2), >670 nm (red fluorescence; FL3), 661 ± 30 nm (red fluorescence; FL4). Samples were run for 5 min at low flow rate (12 ± 3 μl min^−1^) and cytometric data were documented using BD CellQuest Pro software (BD Biosciences). In addition to glaucophyte cells, the green alga *Chlamydomonas reinhardtii* (CC-125), red alga *Porphyridium purpureum* (SAG-1380), cryptophytes *Guillardia theta* (CCMP 2712) and *Chroomonas mesostigmatica* (CCMP 1168), and cyanobacterium *Microcystis aeruginosa* (CPCC 300) were used for comparison. Given the low cell concentration in cultures of *Cyanoptyche gloeocystis* (SAG 4.97) and *Gloeochaete witrockiana* (SAG 46.84), the running time was increased for these taxa to 10 minutes to obtain visible cell clusters. Cytometric surveys of cell mixes and environmental samples included ~ 5,000 APC and PE fluorescent beads (BD CaliBRITE; Becton Dickinson), respectively, as reference. Cytograms prepared with different detector combinations were scrutinized to both identify distinctive emission signals useful for glaucophyte identification and to define narrow cell clusters for further sorting.

**Table 2.**
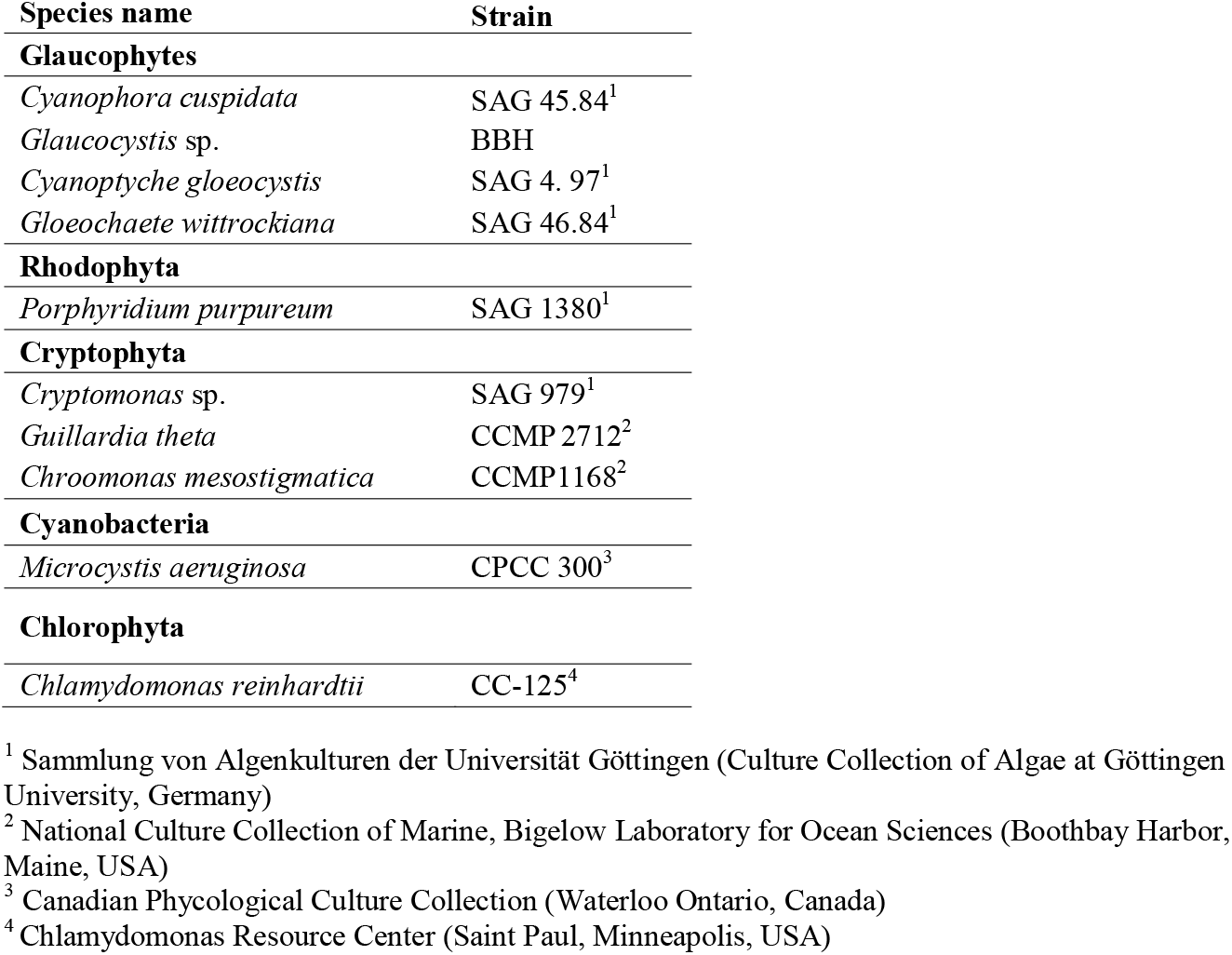
Photosynthetic taxa analyzed.

To comprehensively analyze the glaucophytes signature signal in the red portion of the spectrum, additional strains (Table S1) were analyzed to produce a baseline catalogue of the glaucophyte autofluorescence signature, including species of *Cyanophora* (Fig S1) and *Glaucocystis* (Fig S2).

### Confocal microscopy

To further investigate the autofluorescence profile of glaucophyte cells we performed wavelength (lambda; λ) scanning of *Cyanophora cuspidata* (SAG 45.84) and *Glaucocystis* sp. (strain BBH) live specimens. Five individual cells were identified as regions of interest and excited alternatively with 488 nm and 633 nm lasers (the same and similar excitations wavelengths, respectively, as our FACScalibur instrument), collecting autofluorescence with a 10 nm window for wavelengths between 400 nm to 740 nm. Lambda scans were performed with a Leica TCS-SP2 (Leica Microsystems) confocal microscope.

### Cell counting and viability

Cell recovery efficiency and post-sorting viability were assessed using *C. cuspidata* and *Glaucocystis* sp. BBH cultures. Recovery efficiency was defined as the proportion of cells sorted into the sample tube from a known number of cells (i.e., counted before) that passed through the cytometer fluidics system. For each sample, 1-ml aliquots of exponential-phase cultures were sorted into 50-ml polypropylene tubes using DY-V media as sheath fluid. The cells were recovered in ~50 ml of sheath fluid and centrifuged at 2,500 x *g* for 10 min. The pellet was resuspended in 1 ml of distilled water and enumerated under 40X magnification using a standard nanoplankton chamber (0.1 ml volume). At least 10 random fields of view and/or 200 cells were counted for each subsample (Lund et al. 1958). To facilitate counting of *C. cuspidata* cells, 1-ml subsamples were fixed with 1% formaldehyde, centrifuged and then resuspended in 0.1 to 0.5 ml of distilled water to obtain a reasonable number of cells per field of view.

Additionally, ~50 ml of *C. cuspidata* and *Glaucocystis* sp. BBH cells recovered from identical 1-ml sorting assays were transferred, respectively, into 125-ml sterile Erlenmeyer flasks and incubated in a 14/10 h light/dark cycle at 18 °C to evaluate cell viability and proliferation. An equal number of *C. cuspidata* and *Glaucocystis* sp. BBH control cultures (non-sorted cells) were prepared independently by transferring 1 ml of stock cultures into fresh DY-V media. Cell proliferation was monitored using the same counting procedure described above. Cell culture aliquots of 1 ml were counted every seven days and until a decrease in the number of cells was observed in experimental cultures. Cells densities were used to estimate growth rates for both species following the method of Wood and collaborators (2005).

### Molecular identification

PCR amplifications with primers specific for the plastid 16S rRNA (SSU) locus (16S_F2 [5’ GCATGCAAGCGTTATCCGGAAT 3’] and 16S_R2 [5’ GTTCTTCGCGTTGCATCGAATTA 3’]) (Burja et al. 2001) were carried out to verify the identity of the sorted cells that proliferated during the viability assays. Amplification conditions comprised of a denaturation step at 94 °C for 4 min, then 38 cycles of 94 °C for 1 min, 46–50 °C for 30 s and 72 °C for 2 min, concluding with a 10 min extension at 72 °C. Amplicons were purified using a 5 PRIME PCR Purification kit (now manufactured by Fischer Scientific) and then Sanger sequenced at Genome Quebec Innovation Center.

## Results

### The glaucophyte fluorescence signal

The FL4 detector (661±16 nm) revealed a red signal equal to or greater than the intensity of the APC reference beads (maximum emission, λ=660 nm) in subpopulations of all glaucophytes analyzed (Figures 1 B, E and 2 A, C). This red signal was consistent with the autofluorescence profile of *C. cuspidata* and *Glaucocystis* sp. BBH cells (maximum emission peaks at 668 nm and 675 nm, respectively; Figure 3 B, D) excited with the 633 nm laser of the confocal microscope and overlapped with the maximum emissions of pure CPC (λ=640-655 nm) and APC (λ=655-665 nm) (Bryant 1982; Sonani et al. 2016). This suggested that the glaucophyte signal recorded with the FL4 detectors was caused by the fluorescence of these two blue phycobiliproteins.

**Figure 1.**
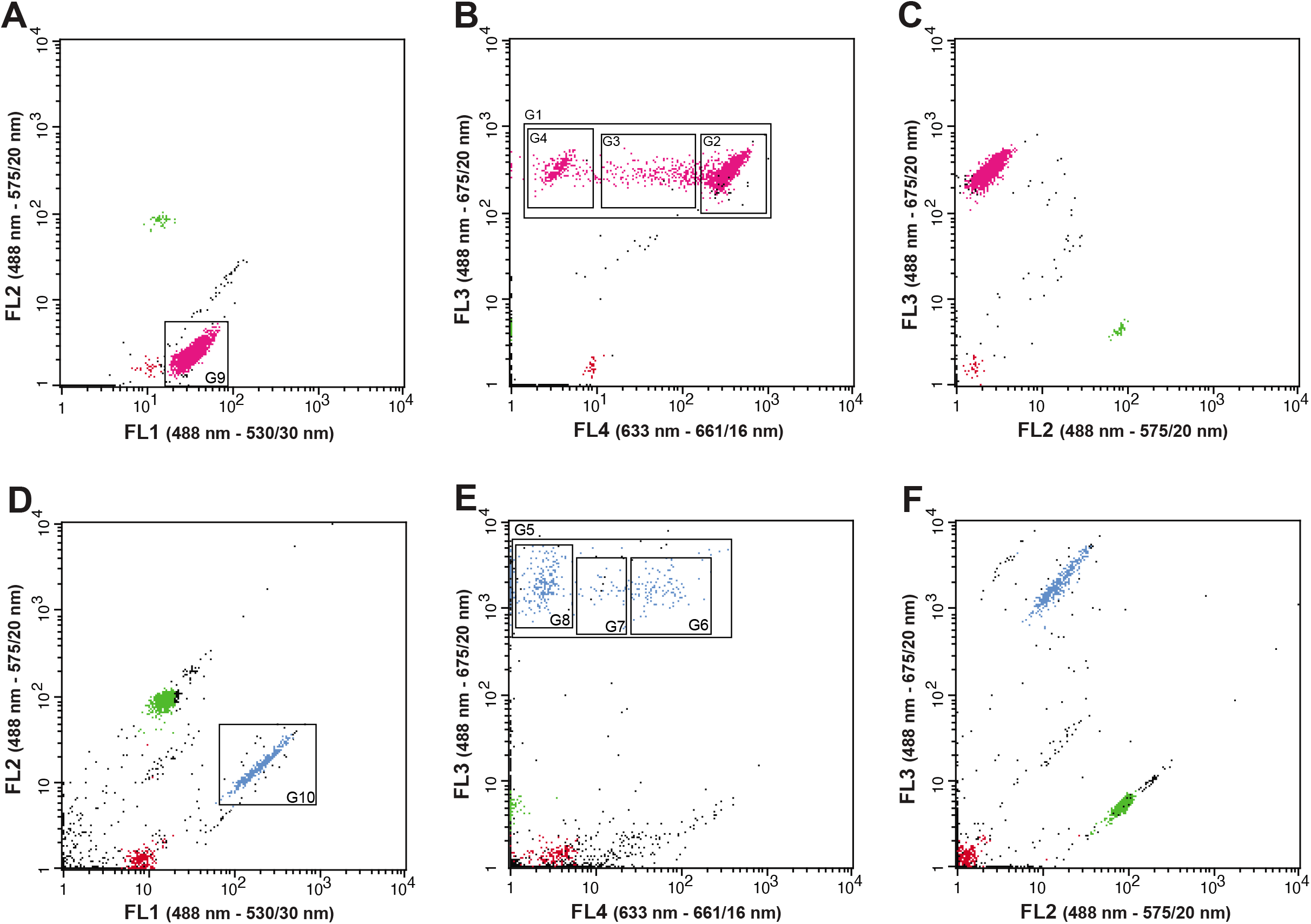
Bi-dimensional cytograms of *Cyanophora cuspidata* SAG 45.84 (**A, B, C**; pink dots) and *Glaucocystis* sp. BBH (**D, E, F**; light blue dots) cultures. Red and light green dots represent allophycocyanin (APC) and phycoerythrin (PE) reference beads, respectively. Different gates (G*N*) used in sorting and viability experiments (see text) are indicated with square areas.

**Figure 2.**
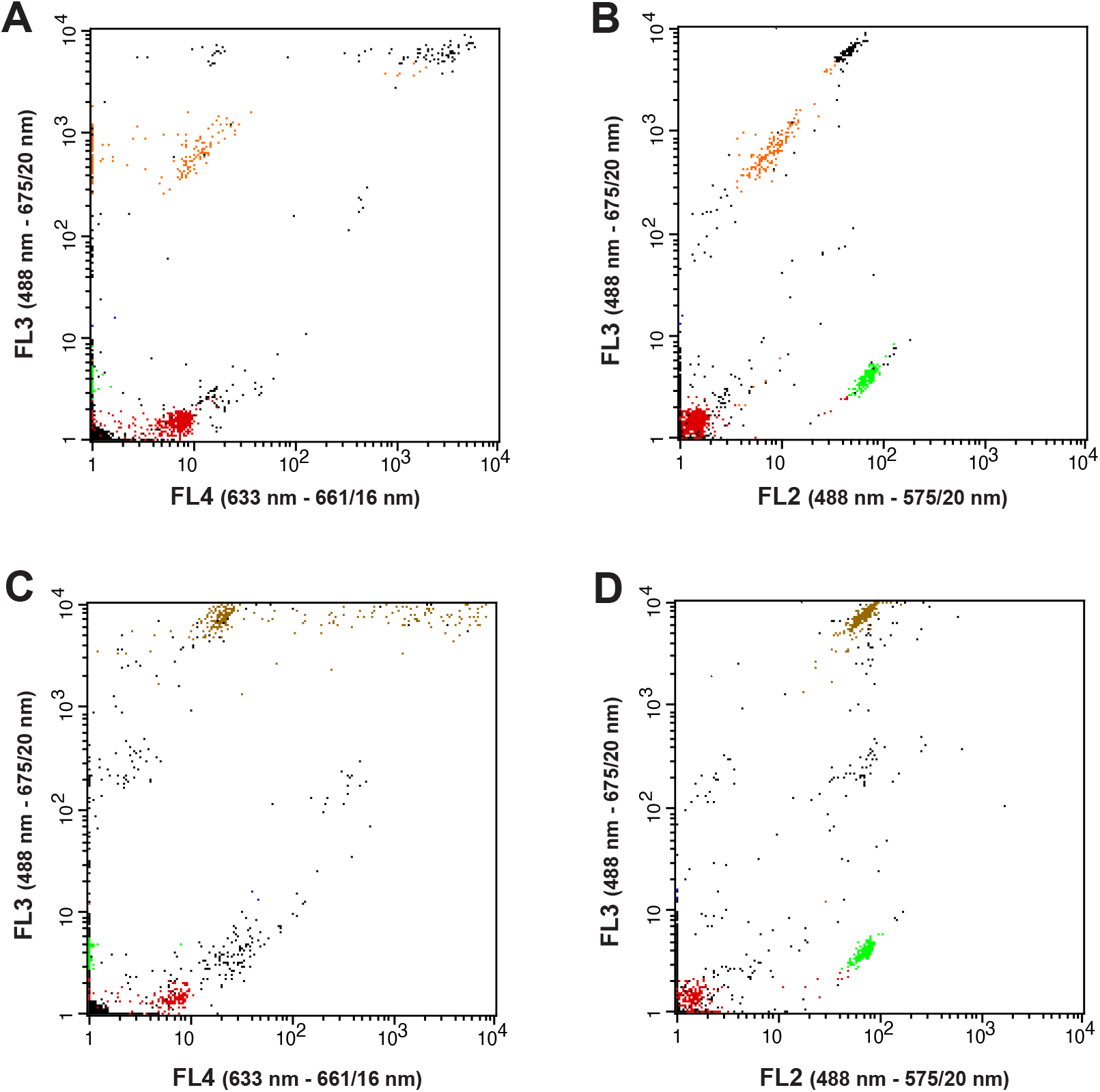
Bi-dimensional cytograms of *Cyanoptyche gloeocystis* SAG 4.97 (**A, B**; orange dots) *Gloeochaete wittrockiana* SAG 46.84 (**C, D**; brown dots). Red and light green dots represent allophycocyanin (APC) and phycoerythrin (PE) reference beads, respectively.

**Figure 3.**
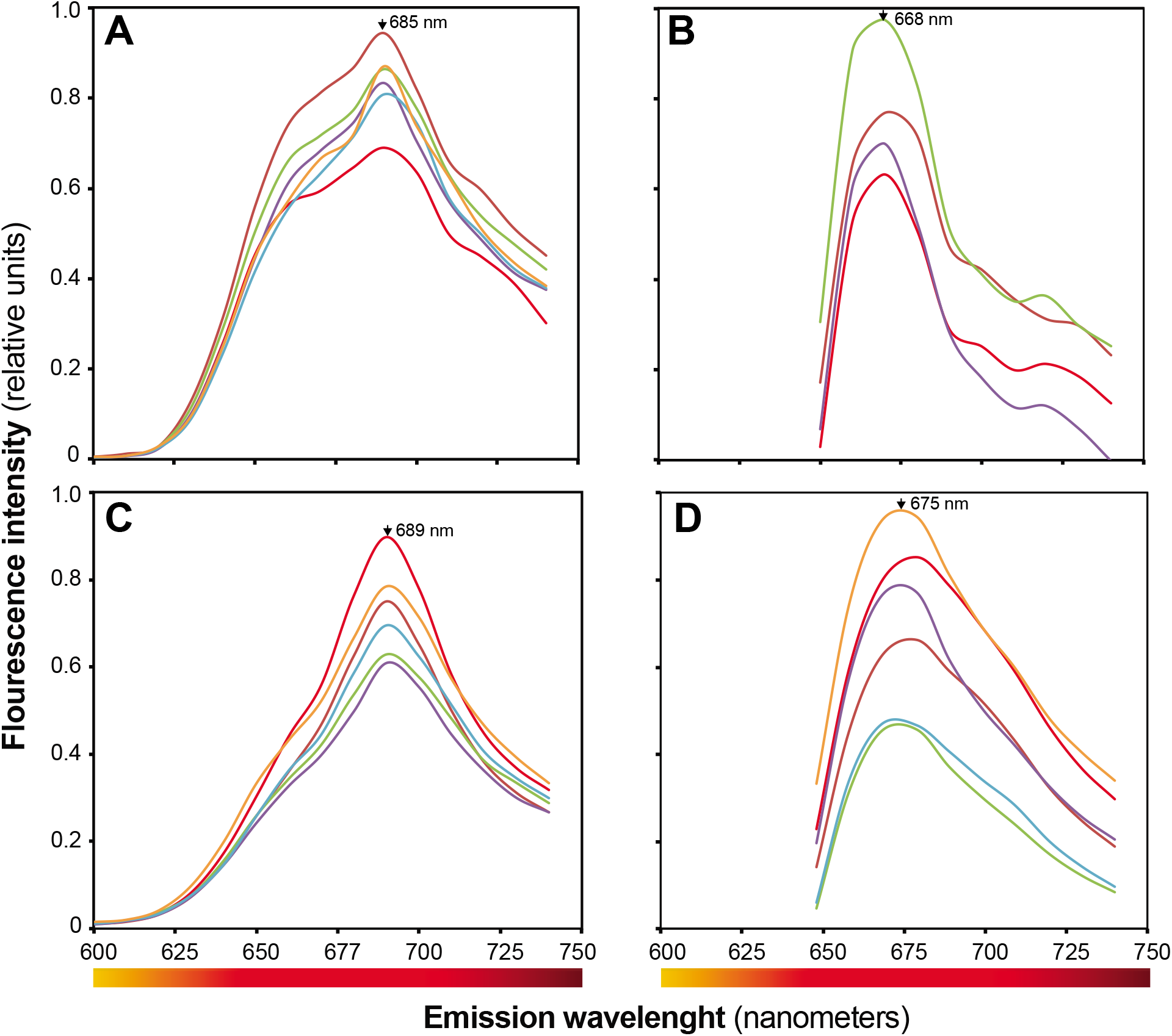
Lambda scan intensity versus emission wavelength (600–750 nm) for *Cyanophora cuspidata* (**A, B**) and *Glaucocystis* BBH (**C, D**) cells. Emission wavelength was recorded using excitation wavelengths of 488 nm (**A, C**) and 633 nm (**B, D**). Maxima emission peaks are indicated with arrows.

In addition, the fluorescence recorded by the FL4 (661±16 nm) FL3 detectors for *C. cuspidata* (Figure 1B) and *Glaucocystis* sp. BBH (Figure 1E) was compared with three other *Cyanophora* species (Supplementary Figure S1) and four *Glaucocystis* species (Supplementary Figure S2). These results produced a consistent set of fluorescence signatures of the group that led to a further step in our identification protocol. We used narrow contiguous sorting gates (Figures 1B, E) and light microscopy to verify that the CPC/APC-positive events scattered along the FL4 detection interval were indeed glaucophyte cells. The distribution of glaucophyte cells along the FL4 range suggested that the phycobiliprotein signal was not homogeneous among the cells in the cultures (Figures 1B, E). In contrast to the dispersed distribution in the FL4, *Cyanophora* and *Glaucocystis* appeared confined in dense clusters (Figures 1A, D) when the FL2 detector (575±20 nm) was paired with green fluorescence (FL1 detector 488-530/30 nm). The low orange signal detected in the FL2 channel (Figures 1C, F) was consistent with the reported absence of PE-like signals in *Cyanophora* and *Glaucocystis* (Chapman 1966). A similar result was obtained also with *Cyanoptyche* (Figure 2B), but in the case of *Gloeochaete* (Figure 2D) some subpopulations presented emissions similar to the PE reference beads (maximum emission, λ=578 nm). To our knowledge, the presence of PE in *Gloeochaete* has not been reported. The cytometric survey also revealed CPC/APC-positive events that correspond to cyanobacteria (corroborated via cell sorting, PCR and DNA sequencing) contaminants in the *Glaucocystis sp. BBH* and *Cyanoptyche gloeocystis* cultures.

### Analyses of photosynthetic cells in mixed cultures

To evaluate the utility of the entire glaucophyte autofluorescence repertoire to delineate exclusive cell-sorting gates, we used the same cytometer settings to examine artificial cell mixes that included glaucophytes and representatives of other photosynthetic lineages known to contain blue and/or red phycobiliproteins (see Table 2 for a complete list of the analyzed groups). Most cell types separated into distinct clusters using both the forward scatter-FL3 (661±30nm) and the FL4-FL3 detector combinations, however some glaucophyte populations overlapped within other cell clusters, precluding simple identification (Figures 4B, C). Signal overlapping occurred in the CPC+APC area (FL4 detector) between certain glaucophyte subpopulations, the cryptophyte *Guillardia theta*, the green alga *Chlamydomonas reinhardtii* and the cyanobacterium *Microcystis aeruginosa* (dotted line semi-ellipses Figure 4C). However, glaucophyte cultures included subpopulations with a higher red fluorescence than most photosynthetic lineages evaluated (FL4; solid-line semi-ellipses in Figure 4C). One important exception is the marine cryptophyte *Chroomonas mesostigmatica* that contains only blue phycobiliproteins but not PE (McKay et al. 1992) and produced a signal that, under our experimental conditions, is indistinguishable from *Cyanophora cuspidata* cells by all cytometrics (Fig. 5). The high orange fluorescence (FL2) of the PE-containing *P. purpureum* and *G. theta* separated these algal types from well-defined glaucophyte clusters and *Chlamydomonas* (Figures 4A, D). Then, the low orange signal of most glaucophytes is an additional characteristic to facilitate discrimination. In the case of cyanobacterial presence, the smaller size (i.e., forward scatter) of unicellular species is a useful discrimination measure (Figure 4B).

**Figure 4.**
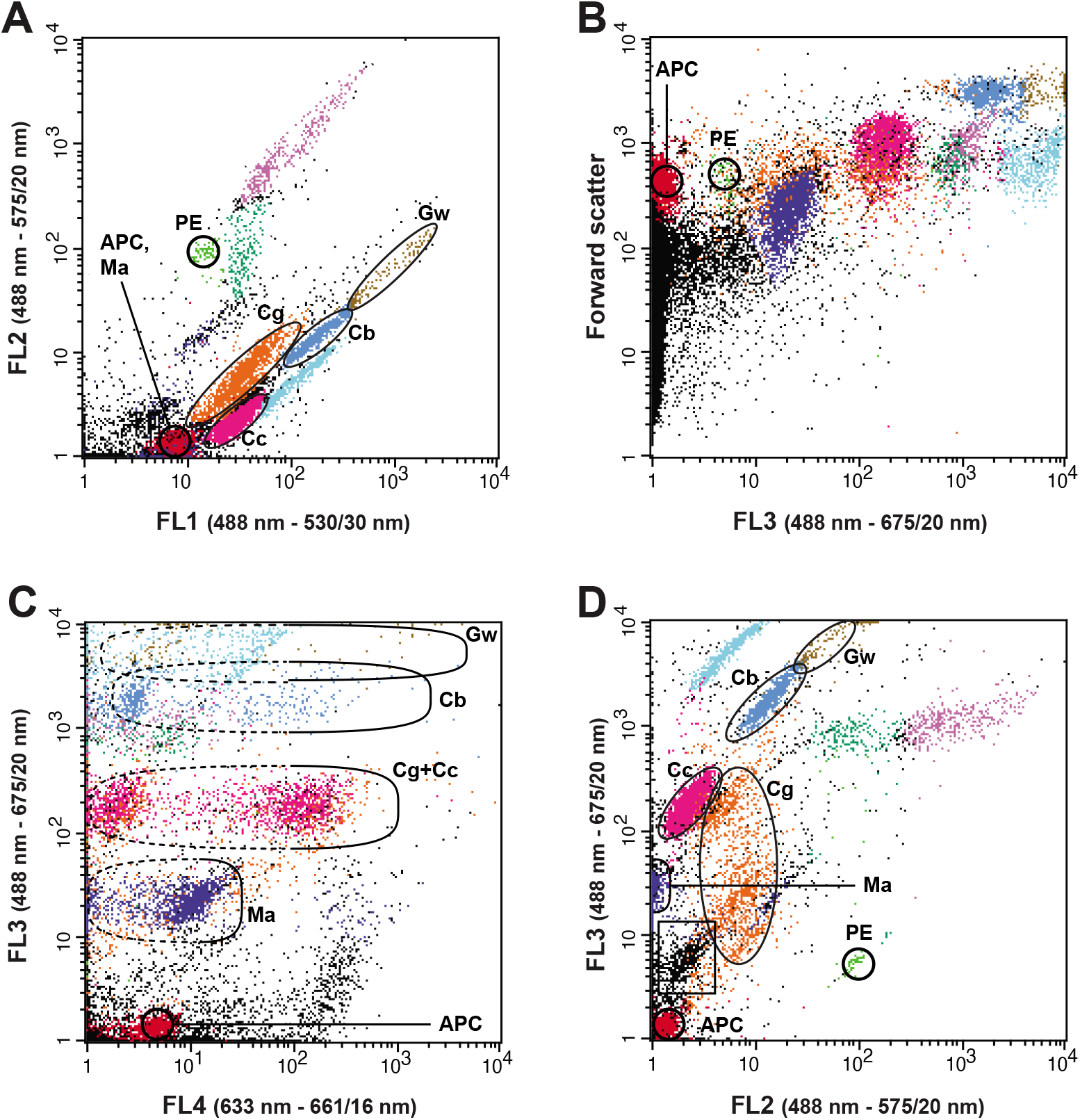
Cytograms of an artificial experimental mix of diverse photosynthetic cell types. The four glaucophyte representatives are circled for visualization purposes. *Cyanophora cuspidata* SAG 45.84 (Cc; pink dots), *Glaucocystis* sp. strain BBH (Cb; medium blue dots), *Cyanoptyche gloeocystis* SAG 4.97 (Cg; orange dots), *Gloeochaete wittrockiana* SAG 46.8 (Gw; brown dots), *Chlamydomonas reinhardtii* CC-125 (light blue dots), *Guillardia theta* CCMP 2712 (dark green dots) *Microcystis aeruginosa* CPCC300 (Ma; dark blue dots) and *Porphyridium purpureum* SAG 1380-1B (purple dots). Allophycocyanin (APC) beads (red dots) and phycoerythrin (PE) beads (light green dots) were included in the analysis. The ellipses delineate the glaucophyte cells dispersed along the FL4 detection range, distinguishing between subpopulations overlapping with other algal types (dotted lines; low relative red fluorescence) and relatively isolated (solid lines; high relative red fluorescence) from other photosynthetic cells in the mix. The black population confined in a square represents a cyanobacterial contaminant coexisting with *Glaucocystis* sp. strain BBH.

**Figure 5.**
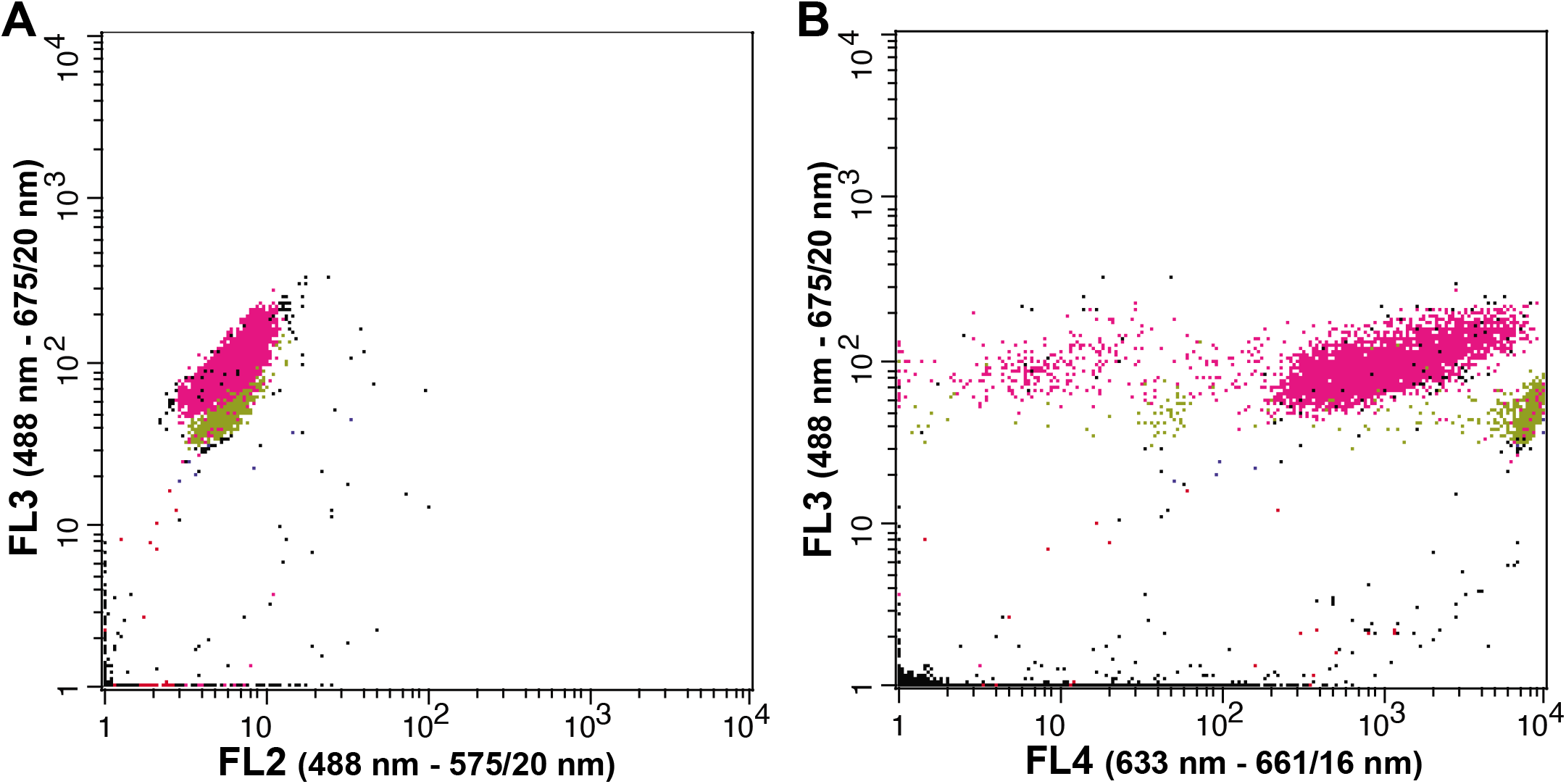
Cytograms obtained from a mix of *Cyanophora cuspidata* SAG 45.84 (pink dots) and *Chroomonas mesostigmatica* CCMP 1168 (light green dots). No reference beads were included in this analysis.

### Cell recovery efficiency

While the high red fluorescence of CPC+APC was one of the criteria we identified to discriminate glaucophytes, the cell populations with positive CPC+APC signals were widely dispersed along the FL4 detection range (Figures 1B, E and 3A, C), which complicates the delimitation of sorting gates. Reducing cell dilution was particularly important for our cell viability and proliferations assays given the slow growth rates of glaucophytes, so we optimized the sorting gates for the recovery of *C. cuspidata* and *Glaucocystis* sp. BBH cells. Wide gates along the FL4 detection range encompassing the majority of the glaucophyte clusters (G1 in Figure 1B and G5 in Figure 1E) produced samples with extremely low cell densities (i.e., below the detection limit of the nanoplankton chamber used). The use of narrower contiguous gates along the same FL4 detection interval (Figures 1B, E) allowed, in the best case, cell recovery efficiencies of 1% for *Glaucocystis* sp. BBH (G8 in Figure 1E) and 1.7% in the case of *C. cuspidata* (G3 in Figure 1B). When narrow sorting gates were defined to contain the denser glaucophyte clusters detected with FL1-FL2 combination (G9 and G10 in Figures 1A, D, respectively), cell recovery efficiency increased to 1.9% for *G Glaucocystis* sp. BBH and 2% for *C. cuspidata*. Based on these results, we selected the gates defined in the green FL1-FL2 combination as the sorting gates for the cell viability/proliferation experiments.

### Viability of sorted cells

Microscopic examination of control cultures and sorted samples indicated some *C. cuspidata* cells had lost flagella (Figures 6A and 7A), likely as consequence of the sorting and centrifugation processes, while *Glaucocystis* sp. BBH cells were morphologically indistinguishable from control cultures maintained in stationary phase (Figures 6B and 7B). Parallel PCR amplification of plastid 16S rRNA sequences from sorted cells confirmed that the sorted cells were *C. cuspidata* and *Glaucocystis* sp. BBH. Intact cells of *Cyanoptyche* and *Gloechaete* were recovered as well, but no post-sorting growth experiments were carried out due to the very low growth rates of these isolates.

**Figure 6.**
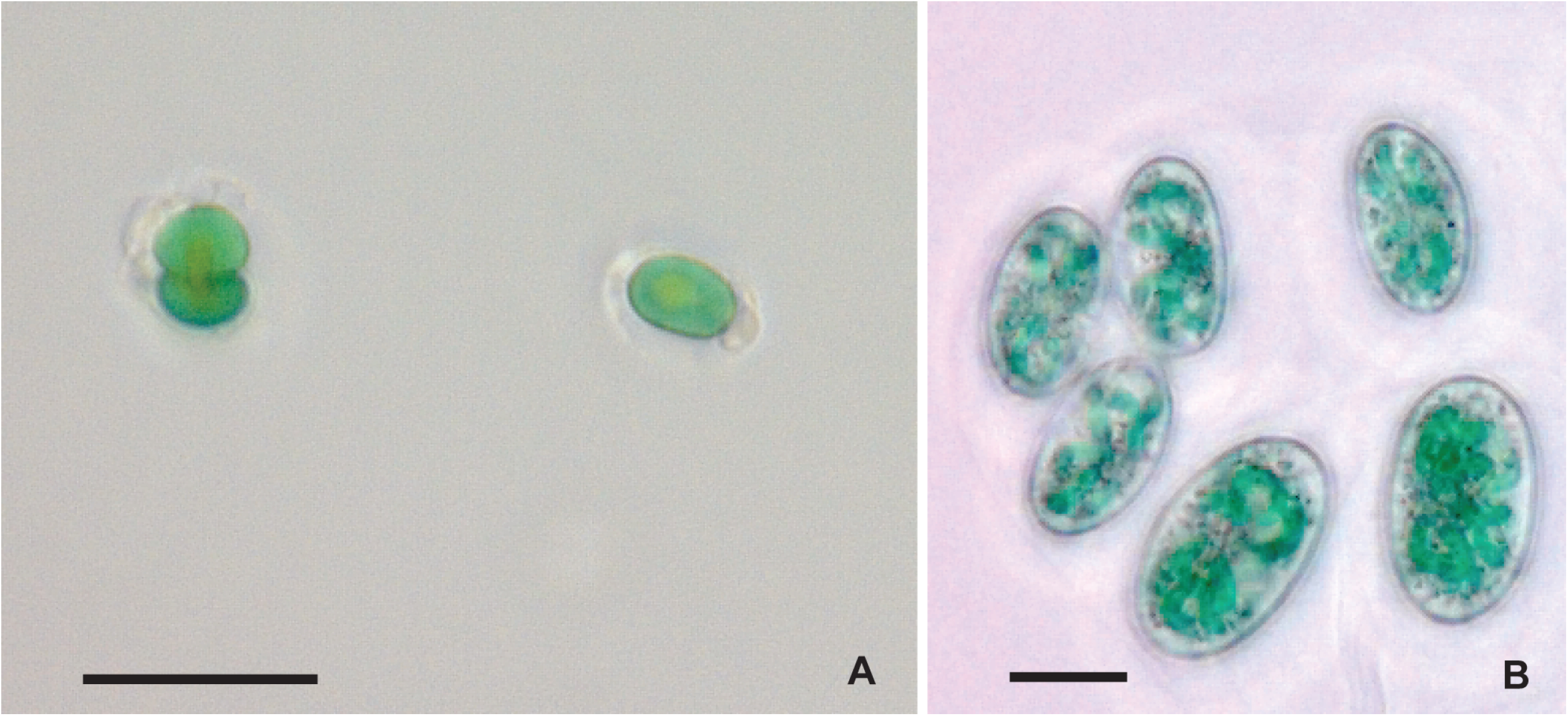
Sorted cells of *Cyanophora cuspidata* (**A**) and *Glaucocystis sp*. BBH (**B**) viewed by light microscopy. No visible cell damage was evident. Scale bars represent 10 μm.

**Figure 7.**
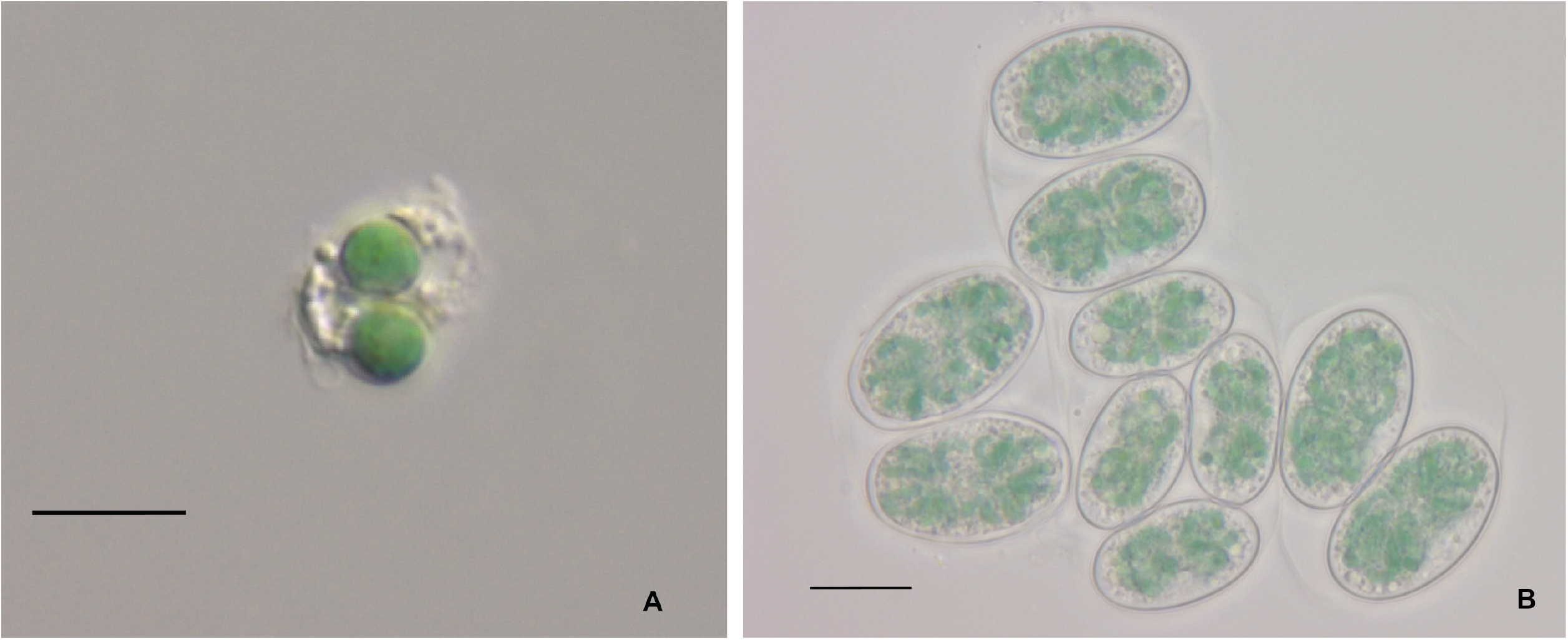
Control culture cells of *Cyanophora cuspidata* (**A**) and *Glaucocystis sp*. BBH (**B**) viewed by light microscopy. Scale bars represent 10 μm.

At least some proportion of cells surviving the sorting protocol grew when incubated in fresh culture media. Post-sorting cell growth of *Cyanophora paradoxa* (strain SAG 29.80) was reported previously, but growth rates were not reported (Sensen et al. 1993). Our *C. cuspidata* control cultures grew at a rate (*r*) of 0.26/day for the first 4 weeks (Figure 8B). Growth in the experimental cultures was undetectable for the first three weeks, but in the following two weeks the sorted cultures exhibited a comparable growth rate (*r=* 0.35/day) to the control cultures (Figure 8A). In the case of *Glaucocystis* sp. BBH, the control cultures presented a growth rate of 0.45/day after 2 weeks (Figure 9 B), whereas the experimental cultures reached rates of 0.36/day after 4 weeks (Figure 9A). To confirm the identity of sorted cells and rule out the possibility of cross-contamination with glaucophyte cells from our own collection, PCR-based identification was performed on sorted cultures six weeks after the sorting procedure. All cultures were confirmed to contain the targeted glaucophyte.

**Figure 8.**
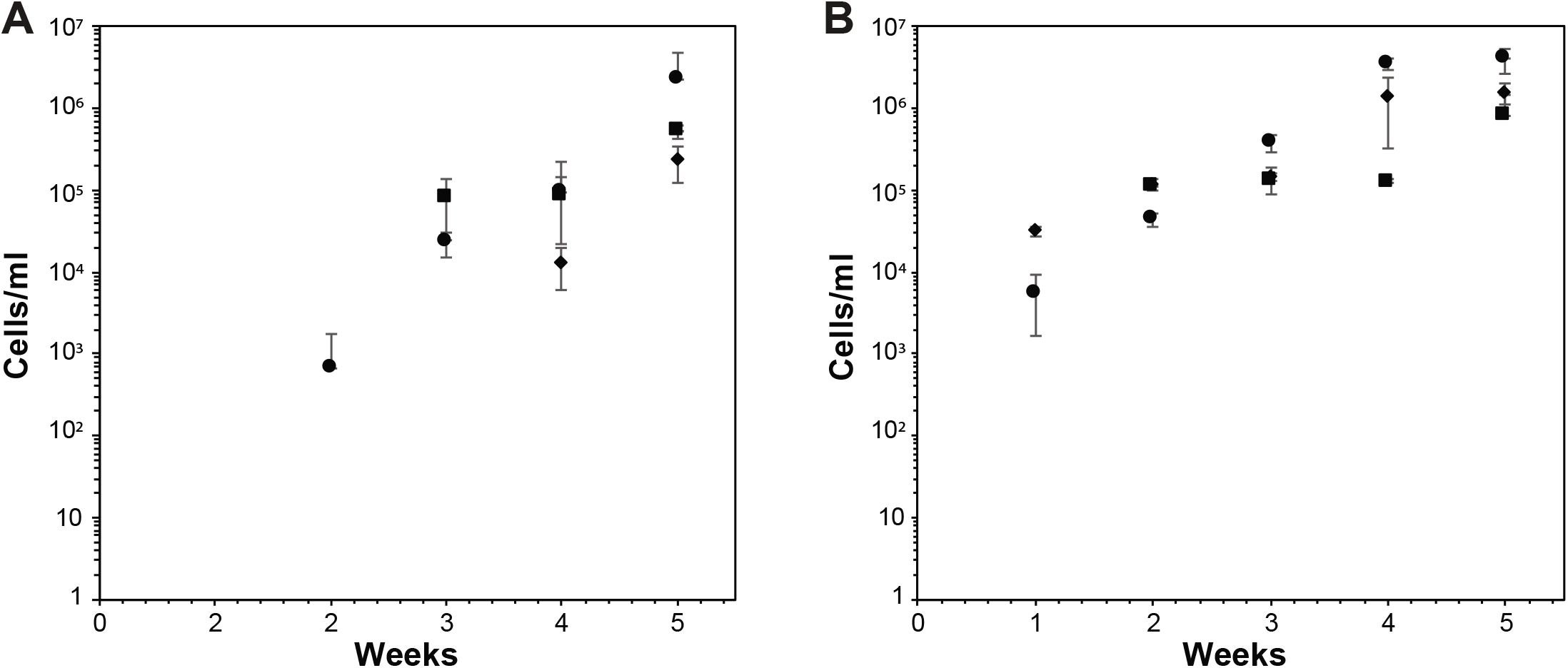
Growth curves of sorted (**A**) and control (**B**) *Cyanophora cuspidata* cells. Each symbol (■, ●, ♦) represents the average of the same starting culture sorted in triplicate (error bars indicate standard deviation).

**Figure 9.**
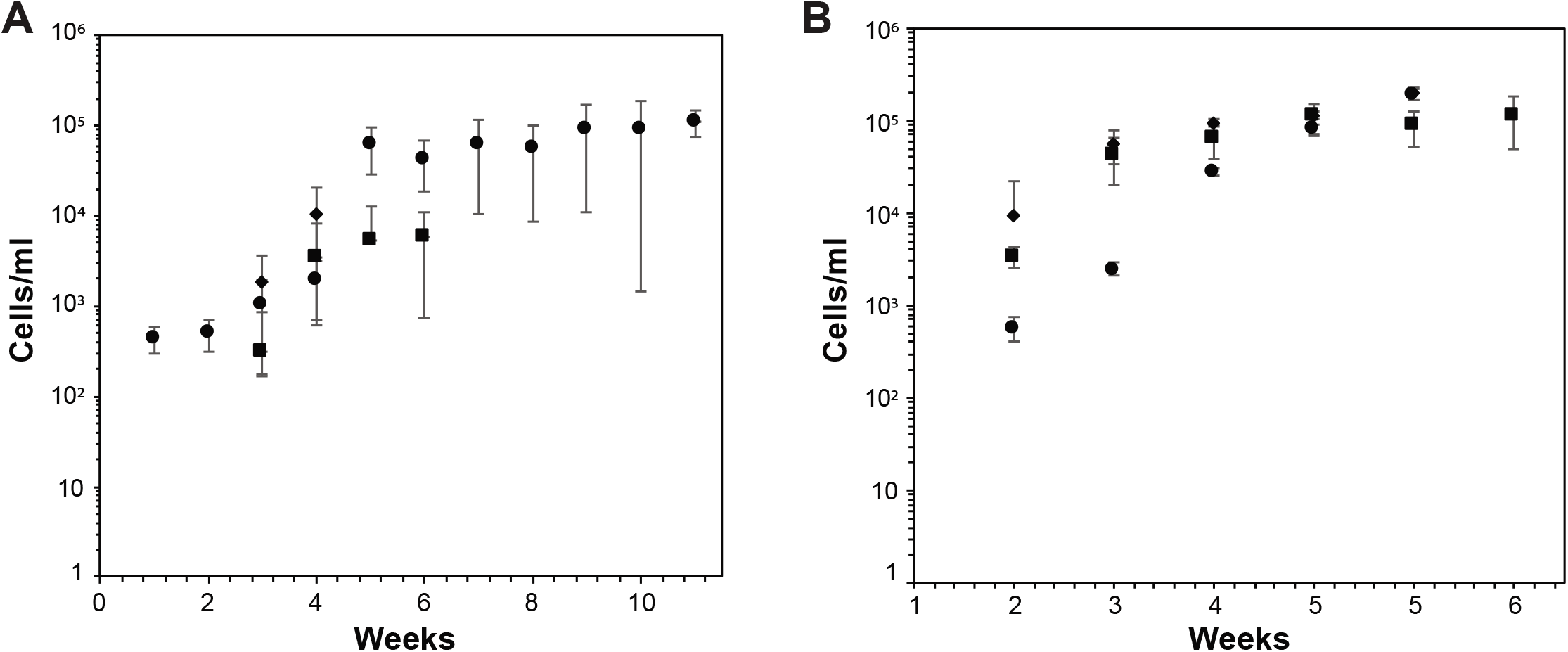
Growth curves of *Glaucocystis sp*. BBH sorted (**A**) and control (**B**) cells. Each marker (■, ●, ♦) represents the average of the same starting culture sorted in triplicate (error bars indicate 1 standard deviation).

## Discussion

### The utility of the glaucophyte autofluorescence signature to isolate viable cells

The cytometric events associated with glaucophyte cells are distributed along the FL4 range, suggesting a non-homogeneous blue phycobiliprotein signal among cells. This is not surprising since metabolic characteristics may change during the cell cycle. Photosynthetic pigment composition, ratio and concentration vary over the different stages of the cell cycle, resulting in the formation of distinct sub-populations in cytograms according to signal intensity (Jorgensen 1966). Nevertheless, the high relative CPC+APC red emission (>660nm) and the low orange signal (i.e., lower than PE reference beads; <587nm) of most glaucophytes investigated are useful traits to detect these cells in complex mixtures of photosynthetic organisms. While our survey did not identify a unique cytometric measure that identifies exclusively glaucophyte cells in complex cell mixes, the comparative analysis indicates that in addition to the relatively high red fluorescence and low orange signal of most glaucophytes, cell size is also useful to distinguish these algae from other photosynthetic cells (e.g., cyanobacteria) in environmental samples. Despite the partial overlapping of the glaucophyte signals with other algal types in certain regions of the red spectra, the high CPC+APC content of glaucophytes facilitates their discrimination from cells with high PE content (e.g., rhodophytes) and PE + blue phycobiliproteins (e.g., the crytophytes *Guillardia* and *Rhodomonas)*. The detection of an orange PE-like signal in *Gloeochaete* is surprising because there are no reports of PE in glaucophytes (Chapman 1966) and no genes encoding PE proteins have been identified in *Cyanophora paradoxa* (Price et al. 2012), neither in transcriptomic data nor plastid genomes of *Cyanoptyche, Glaucocystis* or *Gloeochaete* (Reyes-Prieto et al. 2018). Therefore, it is unlikely that PE is the reason for the *Gloeochaete* orange signal, and further analyses will be required to identify the source. The partial overlap of the glaucophyte cytometric signal with that of other algae may be an obstacle for producing axenic glaucophyte isolates using FACS-based protocols, but the method can be used to produce glaucophyte-enriched samples for molecular identification (e.g., DNA barcoding), which would be a significant improvement over microscopy-manual approaches.

The use of narrow gates (i.e., conservative) increases sorting efficiency and reduces excessive cell dilution in the sorted sample (e.g., Arnold and Lanningam 2010). These considerations were particularly relevant to maximize the utility of the mechanical cell sorter used in our cell proliferation assays. The selection of conservative (e.g., Figure 4A) instead of broad sorting gates (e.g., Figure 4B), allowed us to obtain relatively higher cell densities in sorted samples.

Diverse FACS investigations indicate that not all algal types are able to grow after sorting procedures, limiting the utility of the post-sorting studies to certain cell lineages (Gall et al. 2007; Sinigalliano et al. 2010; Cellamare et al. 2010), but here we were able to recover viable *Cyanophora* and *Glaucocystis* cells using our sorting protocol, establishing a basis for further FACS protocols to establish cultures from novel specimens.

### Development of more efficient sampling strategies

The relatively low efficiency of the mechanical sorting system used in our experiments (300 cells/s for the FACSCalibur) is aggravated when the starting abundance of the target cells is low because sheath liquid constantly flows into the recovery containers even when cells are not sorted (Jochem 2015). In consequence, if the cells are scarce in the original sample (e.g., low glaucophyte density in natural samples), then they will be at a low concentration after sorting, complicating posterior analyses. One option to alleviate this problem is to re-concentrate the cells by centrifugation or filtration after sorting, but these procedures add an additional mechanical stress to the sorted cells. A better option may be to use of high-speed cell sorters (i.e., able to isolate >25,000 cells per second) that will augment the probability of isolating glaucophytes, allowing the separation of hundreds of single cells in individual wells. Single-cell isolation opens the possibility of using culture-independent methods to study glaucophytes. The use of standard molecular identification protocols (e.g., amplification and sequencing of standard genetic markers such as 18S rRNA) immediately after sorting facilitates detection and studies of species diversity.

Another avenue to increase the chances of isolating rare cells is the use of pre-sorting strategies to increase the starting cell density. For instance, tangential flow filtration (TFF) is a method based on a pressurized fluid stream (e.g., environmental water sample) that flows parallel to a filter membrane. While flowing, a fraction of the fluid permeates through the filter and the rest recirculates back to the feed reservoir, progressively concentrating suspended particles (i.e., cells). TFF has been effectively used to increase cell densities of environmental samples for subsequent cytometric analysis (Petruševski et al. 1995; Rossignol et al. 1999, Marie et al. 2010; Balzano et al. 2012). This relatively simple concentration method, paired with FACS protocols, should increase chances of isolating glaucophytes.

Several glaucophytes live in benthic habitats (Takahashi et al. 2016) and most of these taxa form colony-like aggregations under laboratory conditions. The relatively low number of cells recorded from *Cyanoptyche* and *Gloeochaete* cultures is likely caused by cell aggregation, which limits the number of single cells passing through the fluidics system. If similar cell aggregations occur in nature, sampling strategies that maximize the isolation of sessile taxa, such as *Glaucocystis, Cyanoptyche, Gloechaete* and *Chalarodora*, should be considered. For example, mechanical methods to detach cells from leaves or filamentous algae (e.g., shaking and filtration) should facilitate the release of cells into the liquid phase for further analyses (Lawrence et al. 2000). Additionally, physical (e.g., gentle vortexing) or chemical (sample incubation with carbohydrases) procedures to dissociate cells aggregated in mucilaginous compounds may facilitate the release of single cells.

The comparative cytometric analyses, cell sorting and growth experiments described here provide a solid practical framework to guide future investigations of glaucophyte presence, abundance, diversity and geographic distribution. Ultimately, the study of glaucophyte diversity at different levels (e.g., phylogenetic, genomic, metabolic) will enhance our capacity to investigate the origin, evolution and diversification of the eukaryote groups with primary plastids.

## Acknowledgements

We thank John Archibald, Anna Asman (Dalhousie University) and Nicole Poulton (Bigelow Lab for Ocean Sciences) for sharing samples from their culture collections. This work was supported by Natural Sciences and Engineering Research Council of Canada (NSERC) grants to J.L. and A.R.P. A.R.P. is also supported by the Canada Foundation for Innovation (project 28276) and the New Brunswick Innovation Foundation (project RIF2012-006).

## SUPPLEMENTARY INFORMATION

### Supplementary Tables

**Table S1.**
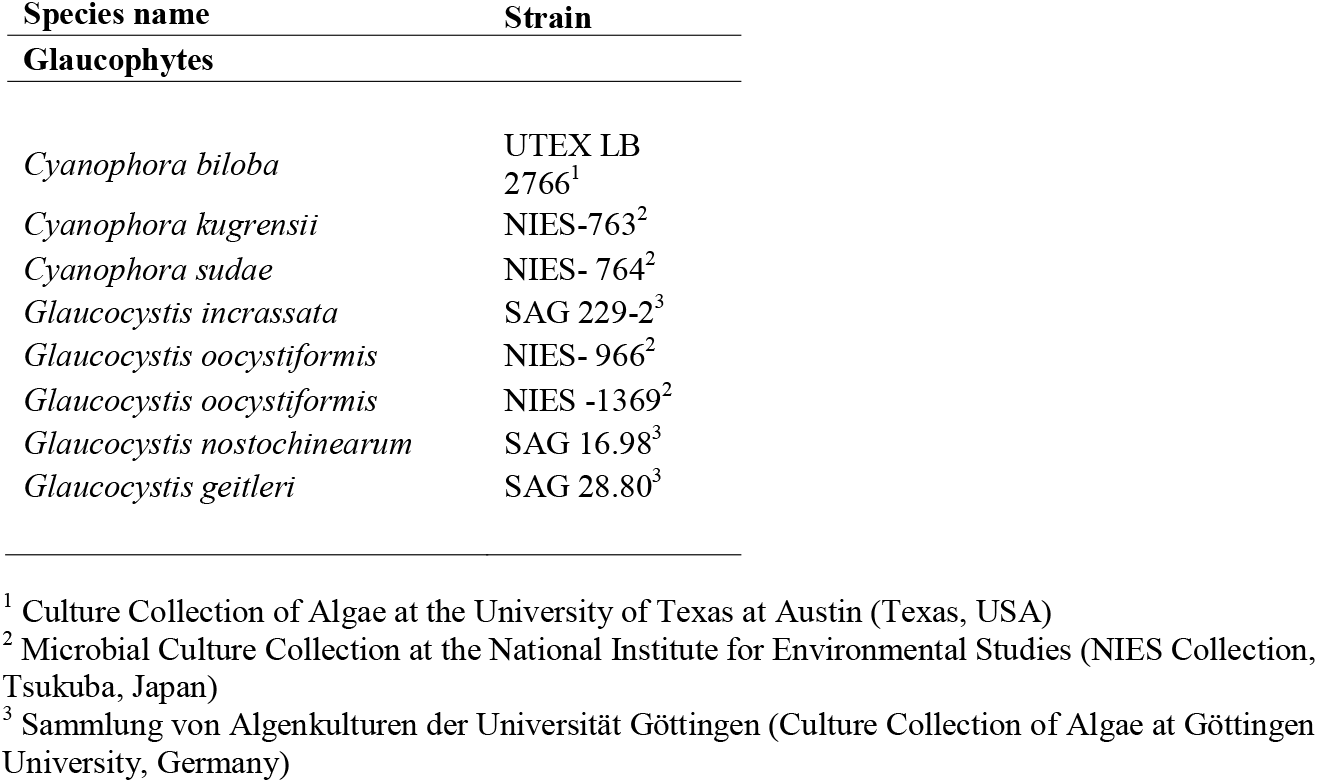
Additional glaucophytes species and strains analyzed.

### Supplementary Figures

**Figure S1**. Cytograms of the *Cyanophora species* analyzed (cell clusters in red), *Cyanophora kugrensii* [(NIES-763) (**A, B**)], *Cyanophora sudae* [(NIES-764) (**C, D**)] and *Cyanophora biloba* [(UTEX LB 2766) (**E, F**)].

**Figure S2**. Cytograms of the *Glaucocystis* species/strains (cell clusters in green). *Glaucocystis incrassata* [(SAG 229-2) (**A, B**)], *Glaucocystis geitleri* [(SAG 28.80) (**C, D**)], *Glaucocystis oocystiformis* [(NIES-1369, NIES-966) (**E, F, G, H**)] and *Glaucocystis nostochinearum* [(SAG 16.98) (**I, J**)].

