## Supplementary Table 1 for "Cytometric Analysis of Diverse Glaucophyte Species Reveals Distinctive Signals Useful for Fluorescence-Based Detection and Sorting"

**Table S1.** Additional glaucophytes strains analyzed

| **Species name** | **Strain** |
| --- | --- |
| **Glaucophytes** |  |
| *Cyanophora biloba* | UTEX LB 2766^1^ |
| *Cyanophora* *kugrensii* | NIES-763^2^ |
| *Cyanophora* *sudae* | NIES- 764^2^ |
| *Glaucocystis incrassata* | SAG 229-2^3^ |
| *Glaucocystis oocystiformis* | NIES- 966^2^ |
| *Glaucocystis oocystiformis* | NIES -1369^2^ |
| *Glaucocystis nostochinearum* | SAG 16.98^3^ |
| *Glaucocystis geitleri* | SAG 28.80^3^ |

^1^ Culture Collection of Algae at the University of Texas at Austin (Texas, USA)

^2^ Microbial Culture Collection at the National Institute for Environmental Studies (NIES Collection, Tsukuba, Japan)

^3^ Sammlung von Algenkulturen der Universität Göttingen (Culture Collection of Algae at Göttingen University, Germany)
