## Supplementary figures and images for "Cytometric Analysis of Diverse Glaucophyte Species Reveals Distinctive Signals Useful for Fluorescence-Based Detection and Sorting"

### Supplementary Figure 1

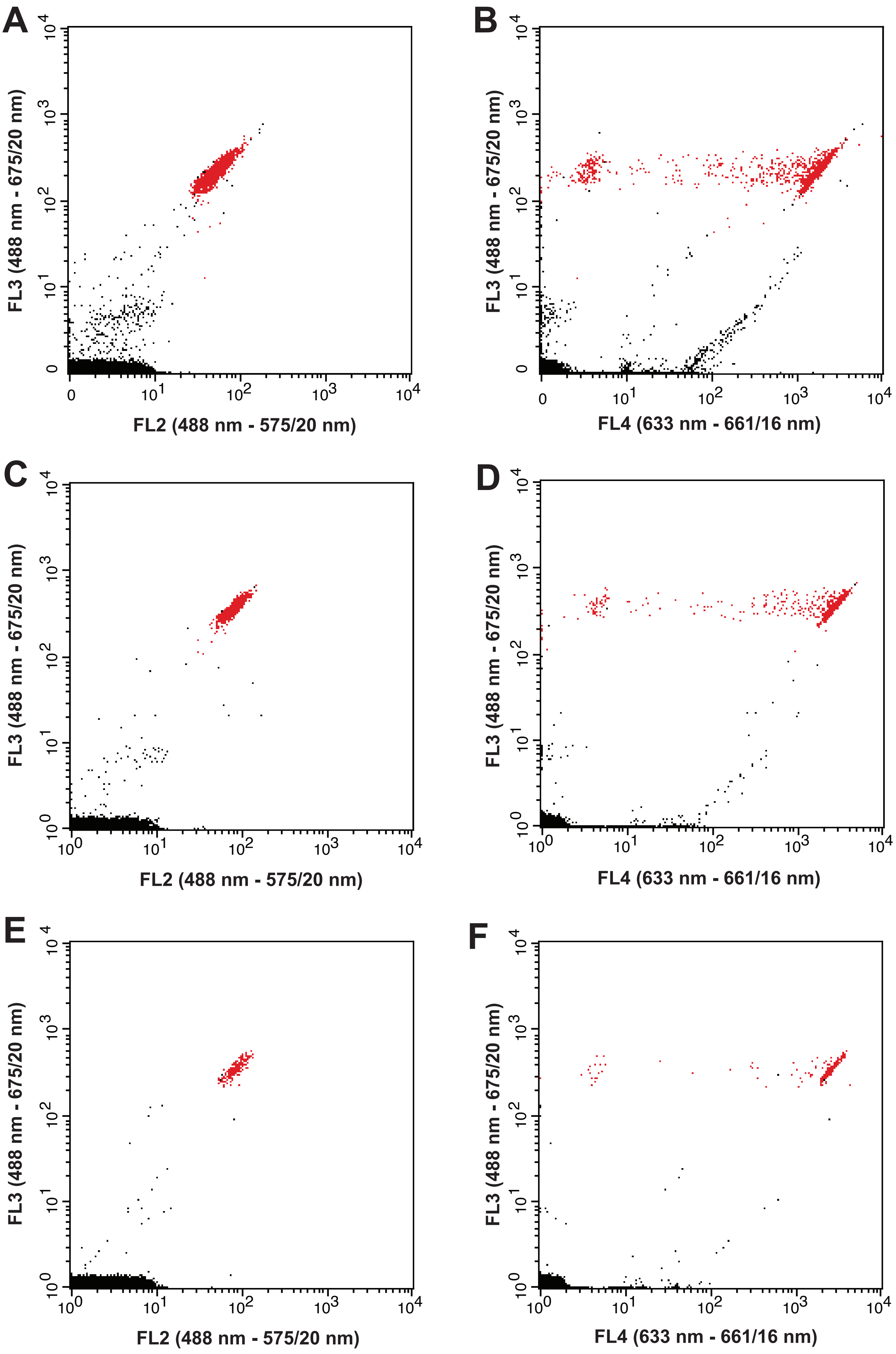

### Supplementary Figure 2

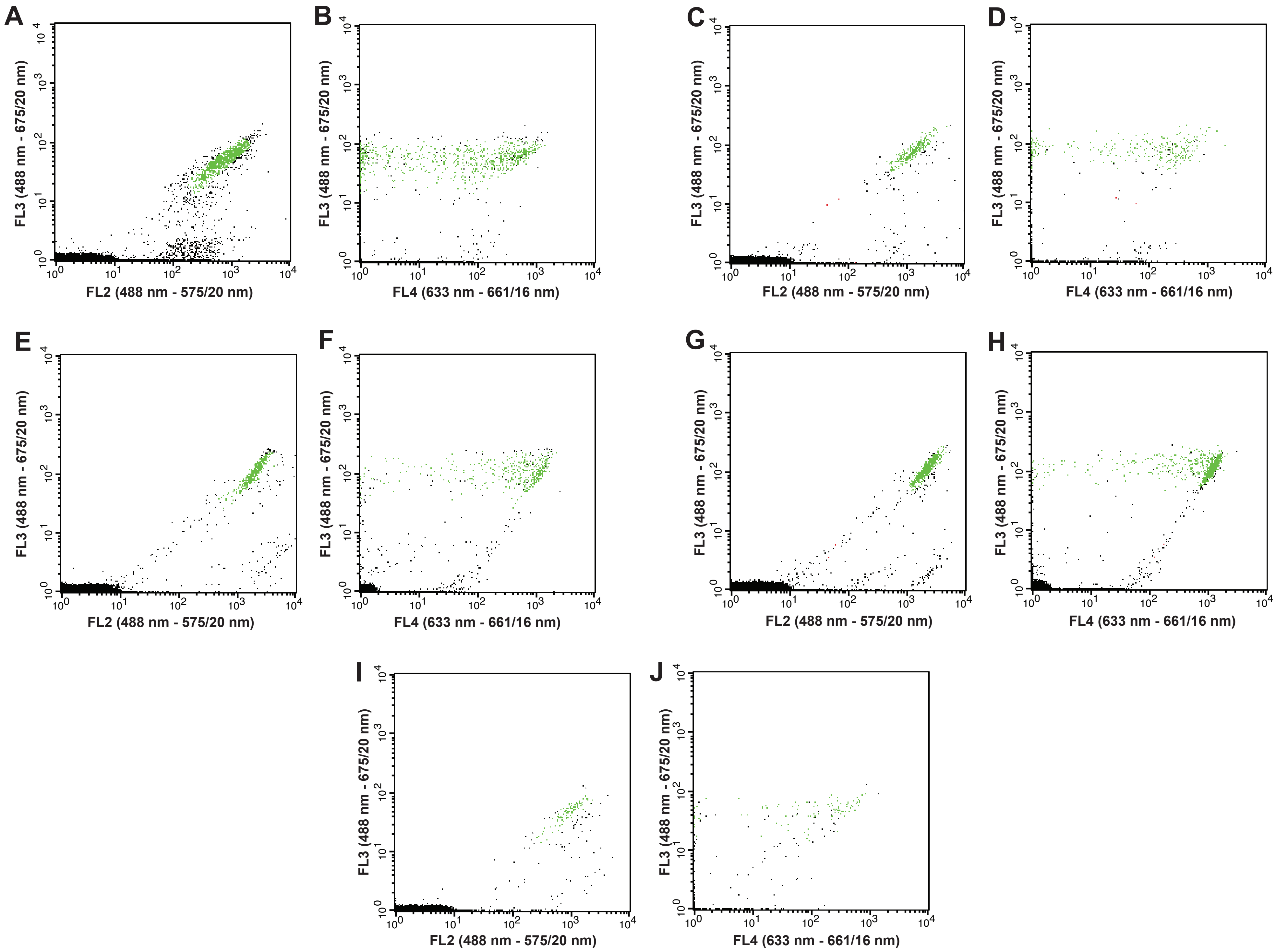
